# Differentiation of semi-transparent tissue phantom inclusions using optical coherence tomography towards label-free neurography and lymphography

**DOI:** 10.1101/2022.05.09.491225

**Authors:** Muhammad Mohsin Qureshi, Taylor Peters, Nader Allam, Valentin Demidov, Alex Vitkin

## Abstract

**Significance:** Lymphatic and peripheral nervous system imaging is of prime importance for monitoring various important pathologic processes including cancer development, metastasis, and response to therapy.

**Aim:** Optical coherence tomography (OCT) is a promising approach for this imaging task but is challenged by the near-transparent nature of these structures. Our aim is to detect and differentiate semi-transparent materials using OCT texture analysis, towards label-free neurography and lymphography.

**Approach:** We have recently demonstrated a novel OCT texture analysis-based approach that used speckle statistics to image lymphatics and nerves *in-vivo* that does not rely on negative contrast. However, these two near-transparent structures could not be differentiated from each other easily in the texture analysis parameter space. Here we perform a rigorous follow-up study to improve upon this differentiation in controlled phantoms mimicking the optical properties of these tissues.

**Results:** The results of the three-parameter Rayleigh distribution fit to the OCT images of six types of tissue-mimicking materials varying in transparency and biophysical properties demonstrate clear differences between them, suggesting routes for improved lymphatics-nerves differentiation.

**Conclusions:** We demonstrate a novel OCT texture analysis based lymphatics-nerves differentiation methodology in tissue-simulating phantoms. Future work will focus on *in-vivo* lymphangiography and neurography studies of longitudinal treatment monitoring for therapy feedback and optimization.

## 1 Introduction

Lymphatic vessels form a network responsible for transporting a colourless, watery fluid called lymph, consisting primarily of proteins and interstitial fluid, from tissue back into the bloodstream^1^. Along with blood vessels, lymphatic vessels play an important role in the metastasis of cancer cells, which is the primary cause of death in cancer patients^2^. The peripheral nervous system can also serve as a conduit for invading cancer cells, facilitating metastasis, and modulate activity and growth of the tumours it innervates^3,4^. In humans, the presence of metastasized tumour cells in lymph nodes is a strong determinant of a poor prognosis, thus emphasizing their importance in oncology^1,2^.

*In-vivo* imaging of lymphatic vessels and nerves is challenging because of the translucent / transparent nature of these structures^5,6^. Typically, lymphatic vessels are detected by the interstitial injection of an exogenous contrast agent or dye which is preferentially absorbed into the lymphatic vessels as they uptake interstitial fluid^7,8^. Further, visualization of lymphatic vessels with such methods is mostly confined to the vicinity of the contrast agent’s injection site. Thus, current gold-standard modalities for *in-vivo* lymphography are far from ideal^8,9^.

Optical coherence tomography (OCT) is a non-invasive, contrast-agent-free, volumetric imaging technique which has already shown promise towards *in-vivo* lymphography^10–12^ and neurography^13,14^. Owing to the above-mentioned transparency of these structures, most OCT approaches so far have relied on ‘negative contrast’ for detection^10,11^, whereby the absence of a signal in a region surrounded by an otherwise signal-rich region indicates a lymphatic vessel. But with increasing imaging depth the SNR drops, and it becomes progressively harder to differentiate absence-of-signal lymphatics from background noise. For instance, several studies have successfully implemented such methods for lymphography using swept-source OCT (ss-OCT) systems only up to a depth of ∼0.5 mm^15^. In our previous studies^12,16^, we presented a novel methodology for visualizing lymphatics (and somewhat unexpectedly also detecting nerves), based on texture analysis of spatial speckle statistics. However, the newly *in-vivo* detected lymphatics and nerves were not clearly differentiated.

In this controlled tissue phantom study, we thus continue developing / refining our innovative methodology to distinguish lymphatics from nerves. We perform a rigorous speckle statistical analysis based on three parameters of the Rayleigh probability distribution function (PDF) in phantoms that mimic different biological structures of interest (tissue-like Intralipid with transparent and near-transparent solid and fluid inclusions). As the first step, we demonstrate the reliable use of the Rayleigh PDFs to differentiate between low-scattering structures (transparent fishing line, plastic tubing, and transparent fluid) and Intralipid tissue-like media, using goodness-of-fit metric as previously shown *in-vivo*^12^. As the novel second step, our analysis then utilizes the Rayleigh PDF’s fit parameters to further differentiate between the low scattering structures themselves. *In-vivo* validation and application to longitudinal preclinical treatment monitoring studies are discussed.

## 2 Methods

A Fourier-domain swept-source OCT system used in this study has been previously described in detail^12^. Briefly, it utilizes a laser (HS2000-HL, Santec, Japan) with a 20 kHz rotating polygon-based tunable filter, with a central wavelength of 1320 nm, a sweep range of 110 nm, and an average output power of 10 mW. The FOV of the volumetric images acquired in this study was 6 mm × 6 mm × ∼1.5 mm in depth, with 24 B-scans captured at each location (such repeats are needed in our analysis to ensure sufficient PDF speckle statistics). Each B-scan is divided in two frames, with 400 A-scan per frame making the inter-frame interval of 25 ms. The distance between two adjacent B-scans was 3.75 μm with 800 A-scans per B-scan, and 1,600 B-scans overall.

The Intralipid gel phantom was composed of water (89% by weight), gelatin (10%, G2500-500G; gel strength 300, Type A, Sigma-Aldrich Co, St. Louis, MO, USA), and Intralipid (1%, Fresenius Kabi Canada Ltd., Richmond Hill, ON, Canada). The resultant optical properties are comparable to porcine skin with a transport mean free path of ∼ 1 mm^17^. Inside the Intralipid phantom, we placed a polytetrafluoroethylene (PTFE) micro-tube (Masterflex RK-06417-11, Cole-Parmer Instrument Company, IL, USA) with an inner diameter of 305 μm and outer diameter of 762 μm, and a 400 μm diameter fishing line (Selizo Clear Fishing Wire) (see Fig. 1(a)). To mimic lymphatic fluid, we used 1 mL water with two added drops of yellow food colouring (Club House, McCormick, London, ON, Canada). The yellow solution was injected by a syringe pump (NE-1000, New Era Pump Systems, Inc., Farmingdale, NY, USA) into the PTFE tubing at a flow rate of 50 μL/min. In addition to the controlled phantom study, we also OCT-imaged *ex-vivo* samples of pork breast to compare the optical properties and speckle statistics of real biological tissues to that of the Intralipid gel.

**Fig. 1.**
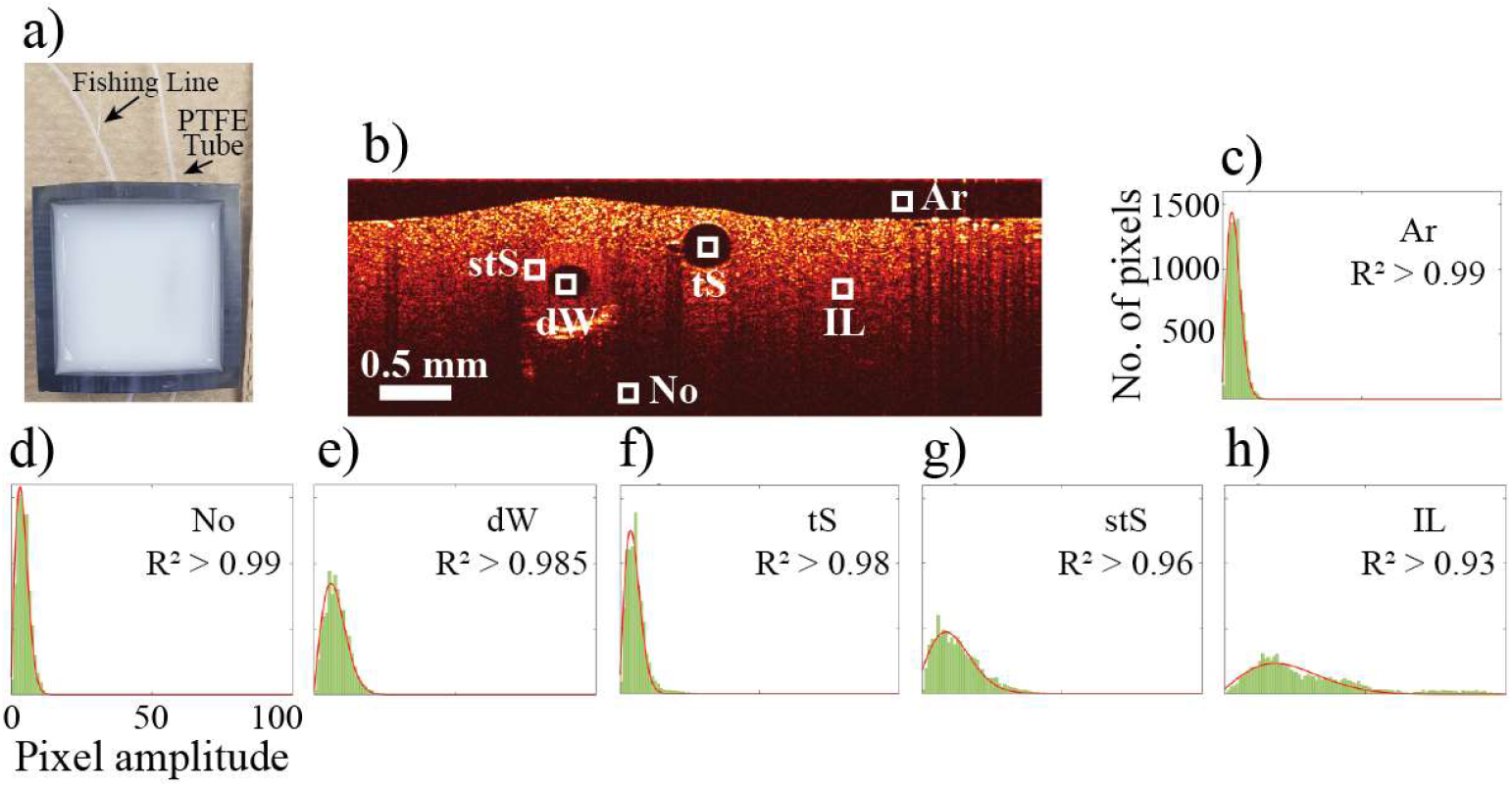
Rayleigh distribution fitting to signal intensity histograms for selected ROIs on an OCT image. (a) Phantom used for this study, showing a fishing line and PTFE tube both embedded in Intralipid-gelatin mixture. (b) A representative B-scan of the phantom, showing six different ROIs for the speckle statistical analysis. (c-h) speckle histograms (green) for the six ROIs: Ar (*R*^2^ > 0.99), No (*R*^2^ > 0.99), dW (*R*^2^ ∼0.985), tS (*R*^2^ ∼0.98), stS (*R*^2^ ∼0.96) and IL (*R*^2^ < 0.93), respectively. The red curves are the corresponding Rayleigh fits. Ar – Air, No – Noise, dW – dyed water, tS – transparent solid, stS – semi-transparent tubing IL – Intralipid

To perform a comprehensive analysis, six regions of interest (ROIs) were identified as air, noise (phantom at depth in the low SNR area), fluid (inside the PTFE tube), transparent solid (fishing line), semi-transparent tubing (PTFE tube wall), and Intralipid (tissue phantom). Each ROI had dimensions of 6 × 10 × 6 (fast lateral × slow lateral × depth) pixels, corresponding to a physical dimension of 45 × 37.5 × 45 μm^3^; with twenty-four repetitive B-scans, this yields a total of a total of 8,640 pixels. For each ROI, pixel intensity distributions were plotted as histograms and fitted with three-parameter Rayliegh PDF^12,18^.

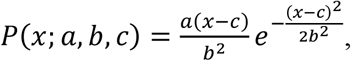

where 𝓍 is the OCT signal intensity. The interpretation of the 3 fitting parameters comes primarily from quantitative ultrasound studies^19,20^ that have been adapted to OCT^12,21^; roughly speaking, *a* is the amplitude normalization parameter, *b* is the scaling parameter, and c is the shifting parameter.

## 3 Results

In order to demonstrate the feasibility of our improved technique to distinguish between lymphatics and nerves, six ROIs were chosen in the tissue-mimicking phantom for the Rayleigh PDF analysis as shown in Fig. 1(b). The resultant signal intensity histograms each contain 8640 pixels (green area in Fig. 1(c)-(h)) with the Rayleigh curve fitting in red. The highest goodness-of-fit range is observed in the ROIs corresponding to noise (No) and air (Ar) (*R*^2^ > 0.99), followed by dyed water (dW) and transparent solid (tS) with 0.99 > *R*^2^ > 0.97, then semi-transparent tubing (stS) with *R*^2^ ∼0.96 and finally Intralipid (IL) with *R*^2^ < 0.93.

Next, we firmed up the fitting statistics by repeating the analysis 20 times. For each ROI, twenty observations were made (*n* = 20) 2 different phantoms and 10 ROIs selected for each of the 6 regions yielding R^2^ values of 0.993 ± 0.0009 (air), 0.994 ± 0.0009 (noise), 0.988 ± 0.0011 (dyed water), 0.983 ± 0.0014 (fishing line / transparent solid), 0.962 ± 0.0135 (PTFE tubing / semi-transparent solid), and 0.910 ± 0.0188 (Intralipid). The results are graphically summarized in Fig. 2. As seen, this approach can clearly differentiate between the Intralipid (∼scattering tissue) and all else (transparent and semi-transparent structures, noise); this is similar to our previous *in-vivo* results^12,22^. However, there is no significant difference between the two transparent / non-scattering media (dyed water and transparent solid); this is again consistent with our previous *in-vivo* result where the lymphatics and nerves could not be distinguished^12^. Therefore, we extend our analysis further, and examine the three parameters of the Rayleigh PDF fits.

**Fig. 2.**
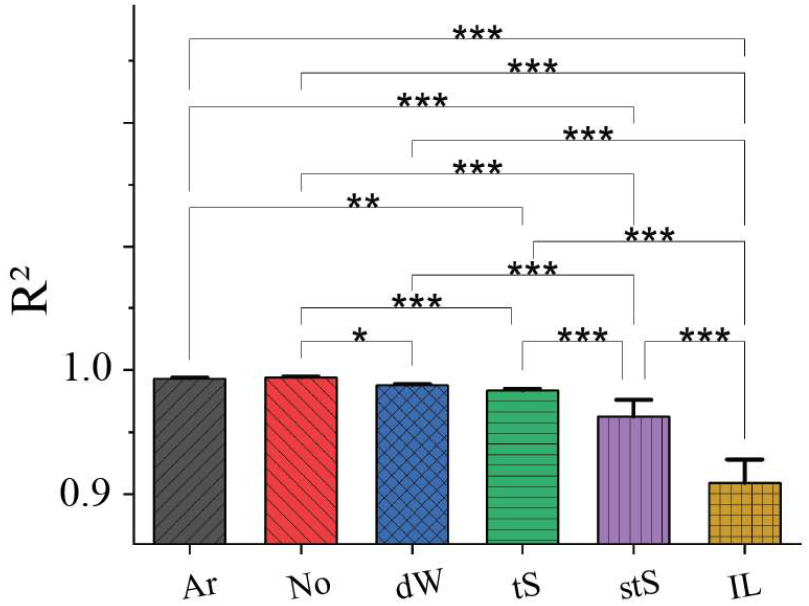
Speckle statistical analysis via Rayleigh fit distinguishes different simulated biological structures. The number of samples for each ROI is twenty (2 different phantoms, 10 representative ROIs selected for each of the 6 regions). (p-values: *p ≤ 0.05; ** p ≤ 0.01; *** p ≤ 0.001)

The results for the ‘a’ and ‘c’ coefficients of the three-parameter Rayleigh PDF for the six ROIs are shown in Fig. 3. The ‘a’ coefficient in Fig. 3(a) is often referred to as the amplitude normalization parameter and appears to be the strongest differentiator between all types of regions: in decreasing order, a = 69.1 ± 5.0 (noise), 41.8 ± 4.2 (air), 41.2 ± 7.5 (transparent solid), 33.1 ± 6.6 (dyed water), 11.2 ± 5.2 (semi-transparent solid), and 3.0 ± 1.7 (Intralipid). This is a noteworthy result as it demonstrates some potential differentiating between the dyed water (a ∼ 33 ± 7) and transparent solid regions (a∼42 ± 7), the phantom analogues of the lymphatics and nerves respectively (more specifically the myelin sheathing of the latter). Fig. 3(b) shows the ‘c’ coefficient results; quantitatively, they are c = -0.22 ± 0.04 (air), -0.20 ± 0.02 (noise), -0.22 ± 0.03 (dyed water), -0.26 ± 0.06 (transparent solid), -0.21 ± 2.70 (semi-transparent solid), 7.56 ± 5.26 (Intralipid). As seen, the only significant difference in c-coefficient values was between Intralipid and the rest of the five materials. Further, the ‘b’ fitting parameter showed minimal discrimination between any of the 6 types of ROIs (results not shown).

**Fig. 3.**
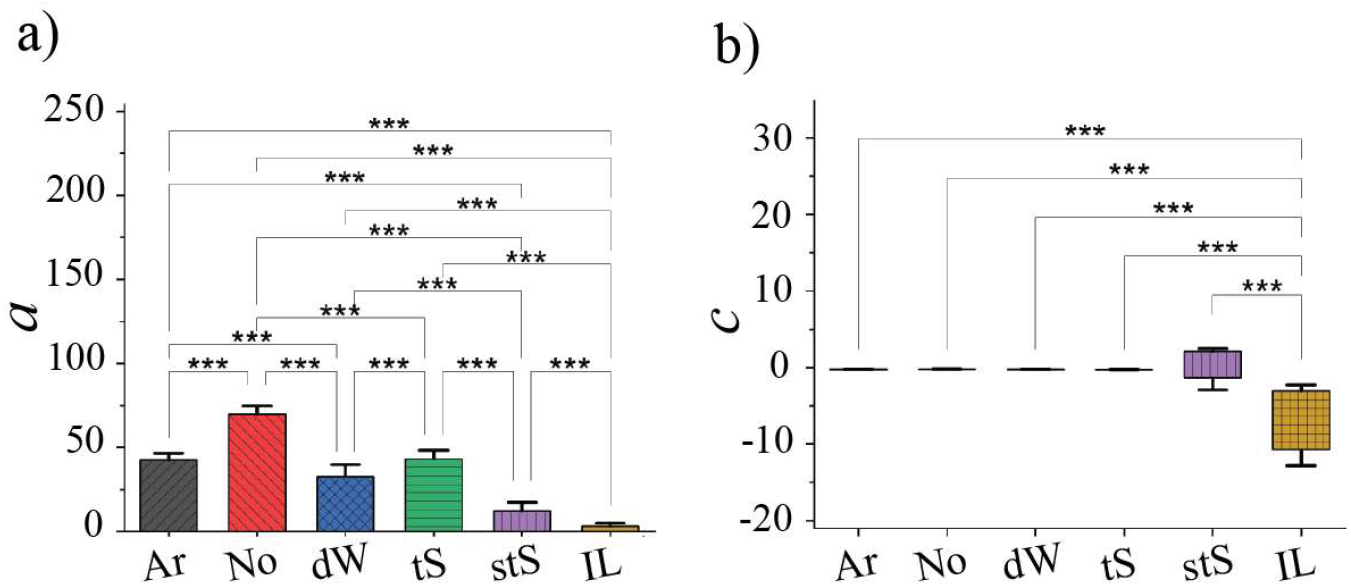
Rayleigh function fit coefficients for the classification of six types of ROIs. (a) ‘a’ and (b) ‘c’ coefficients of the Rayleigh distribution function. A maximum ‘a’ coefficient was observed for the regions of noise. ‘c’ coefficient is significantly different between the Intralipid and the remaining five regions, similar to the R^2^ values trends in Fig. 2. (p-values: *** p ≤ 0.001)

To test the effects of changes in lymphatic flow rates, the water (± yellow food colouring) in the PTFE tubing under different flow conditions was examined. The three experiments involved dyed water flowing at a rate of 50 μL/min, non-flowing dyed water, and non-flowing water (the latter two conditions to focus on the effects of Brownian motion). As shown in Fig. 4(a), the R^2^ values were all very similar (∼0.98 for all three), suggesting that the absence / presence of flow does not affect our Rayleigh fit analysis, at least at the goodness-of-fit level. Looking deeper at the a-b-c fitting parameter space, we find that there are also no significant differences there (representative a-coefficient results shown in Fig. 4(b)). This apparent insensitivity of our methodology to the lymphatic flow conditions may prove useful for *in-vivo* deployment, where this physiological variable is not controlled and in fact varies greatly in inter- and inter-patient setting^23^.

**Fig. 4.**
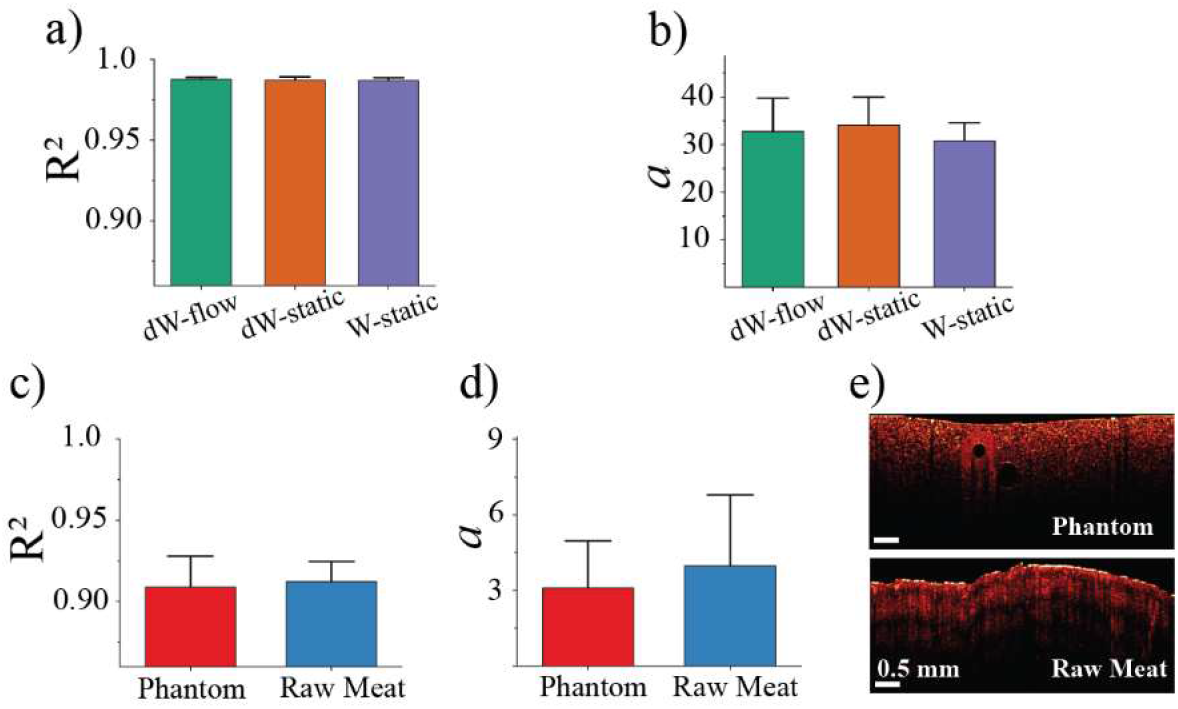
Flow rate effects and phantom-tissue comparison. (a) similarity in the goodness-of-fit R^2^ values and (b) ‘a’ coefficient of Rayleigh distribution function of the fluid ROI, regardless of its flow conditions or presence/absence of dye. (c) quantification of the phantom – biological tissue similarity through R^2^ values and (d) ‘a’ coefficient of the Rayleigh distribution fits. (e) representative B-scans of Intralipid phantom and pork meat, showing similar OCT texture appearance.

To ensure the similarity of the tissue phantom to real tissues in the context of OCT structural appearance, its texture, and its speckle statistics, Fig. 4(c-e) shows the resultant quantitative (panel (c, d)) and qualitative (panel (e)) comparisons. The quantitative analysis via Rayleigh PDF fits summarized in (c) and (d) was performed on 20 different ROIs on phantom and pork meat images, respectively. As seen, the phantom’s optical OCT characteristics are indeed very similar to real tissues in the context of our approach.

## 4 Discussion and Conclusion

In this study we present a method to identify low-scattering structures within a tissue phantom based on OCT texture analysis utilizing speckle statistics. Based on the three-parameter Rayleigh distribution function fit to the pixel intensity distributions within selected image ROIs, this method successfully demonstrated the feasibility to not only detect but also differentiate between different types of transparent and semi-transparent inclusion amid tissue-like scattering background. Importantly, this method differentiates between optically translucent materials, namely solid (fishing line) and fluid (dyes water inside the PTFE tube), which model nerves^24,25^ and the lymph fluid^10,15^, respectively.

For the purpose of this study, axons within nerves can be considered simple fluid filled tubes^26^ where the fluid, similar to cytoplasm^27^, is optically clear (because it is mainly composed of water and thus negligibly scatters light). Fishing wire was selected as a low-scattering object to represent peripheral nerves because it has a low scattering coefficient, being highly transparent due to its homogenous composition^28^ and is both widely accessible and easy to work with. Thus, fishing wire models optical properties of nerves well, for the purpose of establishing an effective detection methodology towards *in-vivo* deployment. For lymphatics that are also difficult to detect due to their optical transparency in the visible and near-infrared spectrum^15^, the underlying cause for their optical transparency is the lack of scatterers (and absorbers) contained in lymphatic fluid; only 6% are solid scattering components (cells, waste products, and/or excess proteins) while the remaining 94% is water^10,15^. Thus, our dyed yellow water with minimal scatterer density is a realistic model for lymphatic fluid. The suitability of our phantoms for simulating the optical properties of nerves and lymphatics provides a useful testbed for optimizing methodology towards eventual *in-vivo* applications.

Imaging of lymphatics and nerves has historically been a challenge, however recent techniques based on OCT imaging are showing some promise^10–13^. This bodes well for detailed preclinical studies, and may have clinical relevance in sites with near-surface pathologies (e.g., skin, epithelial / mucosal lining of numerous body cavities). The typical OCT imaging depth is ∼ 1-3 mm, therefore, the superficially detected lymphatic vessels in human are usually lymphatic capillaries with a diameter range of 10-60 μm^29^ (∼35 μm for mice^30^), similarly at this depth the peripheral nerves in the skin branch directly into the sensory receptors with a diameter of ∼100 – 500 μm^13,28,29^. Leveraging OCT’s limited penetration depth, this work may therefore lead to important applications ranging from early-detection of lymphoedema to compressed nerve diagnosis and surgical guidance.

The long-term objective of this study is to establish OCT as a valuable tool for investigating the full extent of the interactions of the lymphatics and nerves in the tumour microenvironment, which still remains largely unknown today. Due to the important role these semi-transparent structures are known to play in cancer metastasis, this tool may be vital to building a full picture of cancer treatment response (e.g., with radiotherapy) as we work towards adaptive personalized cancer medicine.

### Disclosures

The authors declare that there are no conflicts of interest related to this article.

## Acknowledgments

The OCT system was developed at the National Research Council of Canada with contribution from Drs. Linda Mao, Shoude Chang, Sherif Sherif, and Erroll Murdock.

